# Strategic Decision Making in Biological and Artificial Brains

**DOI:** 10.1101/2025.02.17.638746

**Authors:** Anushka Deshpande

## Abstract

The aim of this paper is twofold. First, it seeks to uncover the algorithms that humans and other animals employ for learning in decision-making strategies within non-zero-sum games, specifically focusing on fully observable iterated prisoner’s dilemma scenarios. Second, it aims to develop a new model to explain strategic decision-making which reflects previous neurobiological findings showing that different brain circuits are responsible for self-referential processing and understanding others. The model stems from the actor-critic framework and incorporates multiple critics to allow for distinct processing of both self and others’ state. We validate the biological plausibility and transferability of our algorithm through comparisons with experimental data from human on the iterated prisoner’s dilemma game.

## 1 Introduction

The emergence of cooperation in human populations has been an object of study in many different disciplines ranging from social sciences, physics, and complex systems to biology and computer science, among many others (Axelrod, 1984). While cooperation is widespread in societies, its origins -remain unclear. While many studies have explored which individual behaviors can promote cooperation in these social dilemmas or aimed to find how much human decision-making aligns with those cooperative strategies, not so much is known about the learning process itself especially when it comes to experimental data where matches don’t end in perfectly cooperative or definitive strategies. As Axelrod(Axelrod, 1984) noted, cooperation does not depend on friendliness but on reciprocity and duration; that is, the stability of cooperation between self-interested agents is learned and stabilizes over time. The field of reinforcement learning (RL) focuses on the theoretical formulation and algorithmic implementation of artificial agents that make sequential decisions to maximize their expected reward (Sutton Barto, 1998) over several iterations through trial and error, which leads to the interesting question -can RL can be an effective tool to understand the evolution of cooperation and understand human decison making in cooperative-competitive settings?

RL has been applied as a theoretical tool for understanding human decision-making behavior (Samejima Doya, 2007; Daw et al., 2006; Kakade Dayan, 2002; Daw, Courville, Tourtezky, 2006; Yoshida Ishii, 2003; Acuna Schrater, 2008; Dayan Daw, 2008). However, in social decision-making, it has not been validated with experimental human behavior as thoroughly as it has been done in partially observable settings (Doshi-Velez Ghahramani, 2001).

Interestingly, in the simple setting of an infinitely repeated prisoner’s dilemma with discounting, randomly initialized RL agents can develop strategies that mimic emergent reciprocity. This raises the question of how humans learn to cooperate and whether RL could be used as a tool to investigate this learning process. As Axelrod noted, cooperation does not depend on friendliness but on reciprocity and duration; that is, the stability of cooperation between self-interested agents is learned and stabilizes over time. In RL, agents often learn over several iterations through trial and error, which makes it an interesting question to ask if RL can be an effective tool to understand the evolution of cooperation.

This paper introduces a novel implementation of the theory of mind (ToM) within a multi-critic actor framework [14], specifically tailored for multiplayer games where full observability is assumed. Inspired by cognitive neuroscience, our model uniquely splits the overarching decision-making policy into two separate tasks: predicting the state of the self and predicting the state of others. Each task is handled by a dedicated critic, mirroring neurobiological findings where different brain areas are responsible for self-referential processing and understanding others. The model’s structure not only facilitates nuanced interactions in game settings by allowing for independent but integrated evaluation of each player’s state but also closely aligns with human cognitive processes as demonstrated in neuroscientific studies.

## 2 Related Work

Previous work in multi-agent reinforcement learning has explored various approaches to strategic decision-making. The DeepPolicy Inference Q-Network (DPIQN) by Hong et al. [7] addresses the challenges of learning effective policies in dynamic environments by using deep neural networks to infer other agents’ policies.

Prashant Doshi’s work on Interactive Partially Observable Markov Decision Processes (I-POMDPs) [4] introduces a recursive modeling approach for multi-agent reasoning. In the realm of inverse reinforcement learning, Ng and Russell’s work [15] on inferring reward functions from observed behavior has been influential. Foerster et al.’s Learning with Opponent-Learning Awareness (LOLA) [5] improves agent performance in competitive environments by considering opponents’ learning dynamics. Our approach shares this goal but achieves it through a novel multi-critic architecture rather than direct opponent modeling.

Wang et al.’s research on modeling mistrust in multi-agent interactions [23] provides valuable insights into trust dynamics. While these prior works have made significant strides in multi-agent learning and decision-making, our study aims to bridge the gap between artificial and biological decision-making processes. We compare human and animal decision-making to common RL frameworks such as DQN [11], PPO [17], and REINFORCE [24], as well as our novel ToM-AC model. Our approach extends the insights from Daw et al.’s study on separable behavioral processes in reward learning [3] by incorporating distinct neural pathways for self and other processing. Furthermore, our ToM-AC architecture, inspired by the feature-specific prediction error model proposed by Lee et al. [10], not only provides a better fit for behavior across species but also achieves superior performance in two-player games. This approach allows us to create a more biologically plausible framework that captures the nuances of decision-making in social contexts more accurately than previous approaches, while also advancing the field of multi-agent reinforcement learning.

## 3 Background

### 3.1 Reinforcement Learning

Reinforcement learning (RL) is a technique for an agent to learn which action to take in each of the possible states of an environment *E*. The goal of the agent is to maximize its accumulated long-term rewards over discrete time steps [21]. The environment *E* is usually formulated as a Markov decision process (MDP), represented as a 5-tuple (*S, A, T, R, γ*). At each timestep, the agent observes a state *s* ∈ *S*, where *S* is the state space of *E*. It then performs an action *a* from the action space *A*, receives a real-valued scalar reward *r* from *E*, and moves to the next state *s*^*′*^ ∈ *S*. The agent’s behavior is defined by a policy *π*, which specifies the selection probabilities over actions for each state. The reward *r* and the next state *s*^*′*^ can be derived by *r* = *R*(*s, a, s*^*′*^) and *T* (*s*^*′*^|*s, a*) = *Pr*(*s*^*′*^|*s, a*), where *R* and *T* are the reward function and the transition probability function, respectively. Both *R* and *T* are determined by *E*. The goal of the RL agent is to find a policy *π* which maximizes the expected return *G*_*t*_, which is the discounted sum of rewards given by 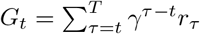, where *T* is the time step when an episode ends, *t* denotes the current time step, *γ* ∈ [0, 1] is the discount factor, and *r*_*τ*_ is the reward received at timestep *τ*. The action-value function (abbreviated as Q-function) of a given policy *π* is defined as the expected return starting from a state-action pair (*s, a*), expressed as *Q*^*π*^(*s, a*) = E [*G*_*t*_|*s*_*t*_ = *s, a*_*t*_ = *a, π*].

### 3.2 Biological Background

The mammalian brain has multiple learning subsystems. Major learning components include the neocortex, the hippocampal formation (explicit memory storage system), the cerebellum (adaptive control system), and the basal ganglia (reinforcement learning, also known as instrumental conditioning).

The frontal dopaminergic input arises in a part of the basal ganglia called ventral tegmental area (VTA) and the substantia nigra (SN). The signal generated by dopaminergic (DA) neurons resembles the effective reinforcement signal of temporal difference (TD) learning algorithms [19, 21]. Another important part of the basal ganglia is the striatum. This structure is made of two parts, the matriosome and the striosome. Both receive input from the cortex (mostly frontal) and from the DA neurons, but the striosome projects principally to DA neurons in VTA and SN. The striosome is hypothesized to act as a reward predictor, allowing the DA signal to compute the difference between the expected and received reward. The matriosome projects back to the frontal lobe (for example, to the motor cortex). Its hypothesized role is therefore in action selection [1, 19, 20].

#### 3.3 Actor-Critic Algorithm

The actor-critic model of the basal ganglia developed here is derived from [1]. It is very similar to the basal ganglia model in [19] which has been used to simulate neurophysiological data recorded while monkeys were learning a task [20]. All units are linear weighted sums of activity from the previous layers. The actor units behave under a winner-take-all rule.

In actor-critic algorithms, the policy structure (actor) and the value function (critic) are separated. The actor updates the policy parameters *θ* in the direction suggested by the critic, which evaluates the action taken by the actor using the value function parameters *ω*. The actor selects actions according to a policy *π*_*θ*_(*a*|*s*), while the critic estimates the value function *V* ^*π*^(*s,ω*), which is used to critique the actor’s actions.

The policy gradient method is used to update the actor’s policy:

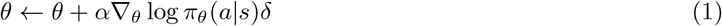

where *δ* = *r* + *γV* (*s*^*′*^, *ω*) −*V* (*s, ω*) is the temporal difference (TD) error, and *α* is the learning rate.

The critic updates the value function parameters *ω* by minimizing the mean squared error of the TD error:

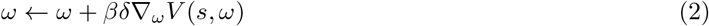

where *β* is the learning rate for the critic.

subsectionGame Theory and Payoff Matrices

Game theory studies strategic interactions among rational decision-makers. In two-player games, payoff matrices represent possible outcomes based on players’ actions. Each cell (x, y) denotes payoffs for Player 1 (x) and Player 2 (y) for a given action combination.

### 3.4 Prisoner’s Dilemma

The Prisoner’s Dilemma exemplifies a conflict between individual and collective interests. The payoff matrix is structured as follows:

Where *T > R > P > S*, and 2*R > T* + *S*. This structure creates a scenario where:

- Mutual cooperation (*R, R*) yields the best collective outcome.
- Defection is individually rational, as *T > R* and *P > S*.
- The Nash Equilibrium is mutual defection (*P, P*), which is Pareto inefficient.

This dilemma highlights the tension between individual rationality and collective optimality, a key focus in multi-agent reinforcement learning research.

This dilemma highlights the tension between individual rationality and collective optimality, a key focus in multi-agent reinforcement learning research.

## 4 Algorithm Description

The Theory of Mind Actor-Critic (TOMAC) is a novel reinforcement learning algorithm inspired by neuroscientific findings on human cognitive processes.

TOMAC operates on pre-decomposed value functions, taking as input separate representations of ‘self’ and ‘other’ values, *Q*_*self*_ and *Q*_*other*_. Its primary function is to effectively utilize and integrate these components through learnable weights. This approach enables adaptive decision-making by dynamically adjusting the importance of self vs. other information based on environmental demands. By separately processing these pre-decomposed streams, TOMAC can develop policies that balance individual and collective interests in complex multi-agent scenarios.

In our multi-agent reinforcement learning framework, we employ a dual-critic architecture comprising a Self Critic and an Other Critic for each agent. These critics estimate action-value functions, with the Self Critic focusing on the agent’s own actions and outcomes, while the Other Critic evaluates interactions and consequences related to other agents’ actions. The update rules for agent *i*’s Self Critic and Other Critic are given by:

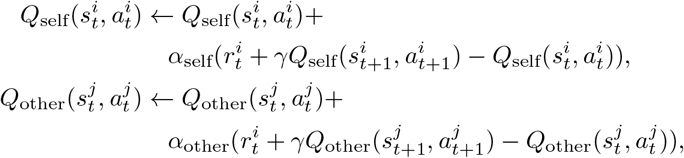

where *α*_self_ and *α*_other_ are learning rates, and *γ* is the discount factor. The actor updates its policy based on a weighted sum of both critics’ outputs:

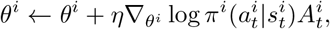

where 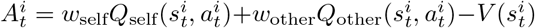 is the advantage function. Here, 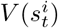 represents the state value function, mathematically defined as:

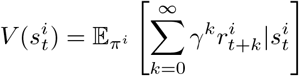

which estimates the expected cumulative discounted reward from state 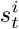 following policy *π*^*i*^. The weights *w*_self_ and *w*_other_ are learnable parameters that balance the influence of self-centered and other-aware critics in the policy update.

### Algorithm 1

Algorithm: Theory of Mind Actor-Critic (TOMAC)

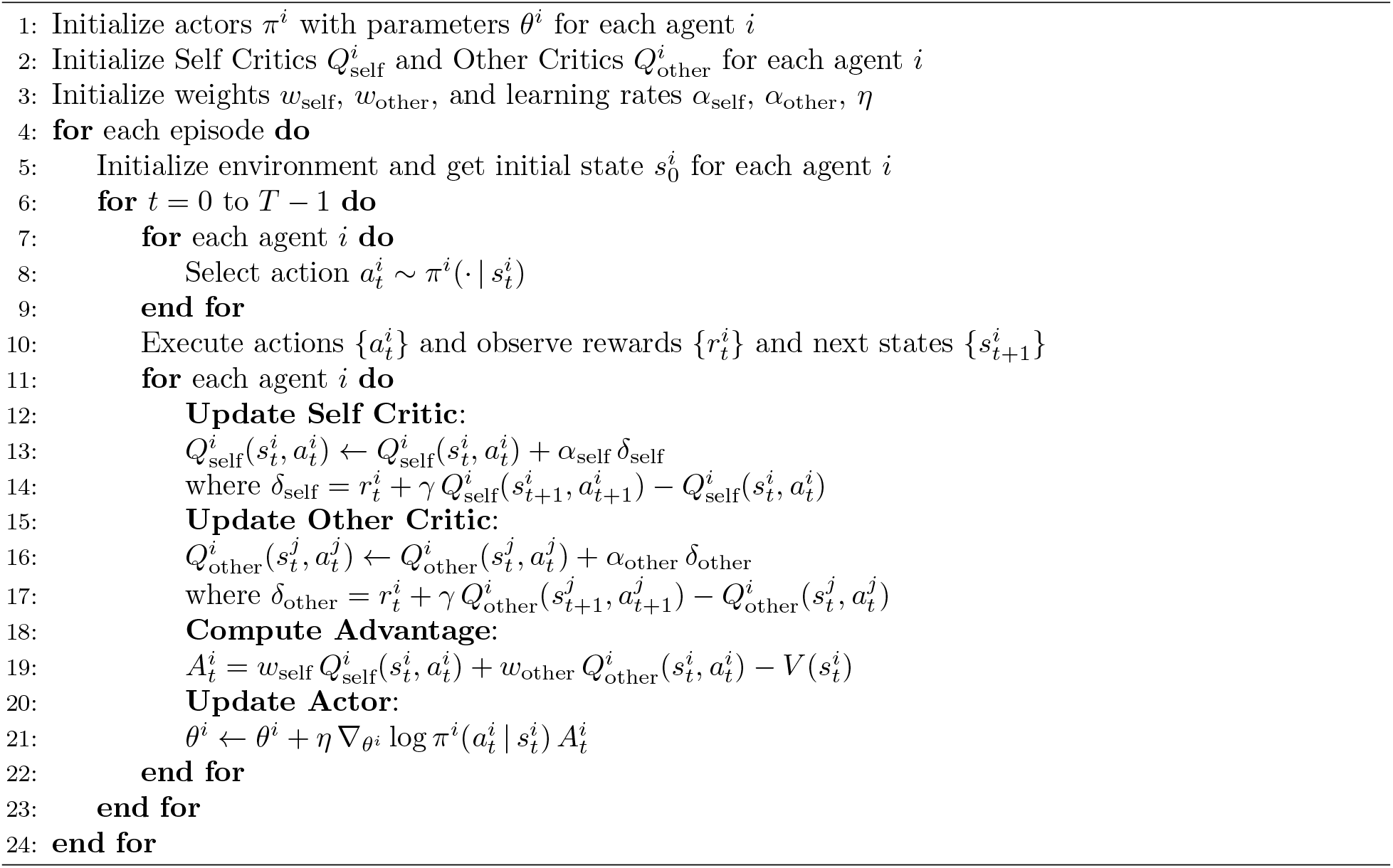

In this algorithm, each agent *i* maintains its own actor *π*^*i*^, Self Critic 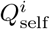, and Other Critic 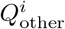. The learning process occurs simultaneously for all agents, allowing them to adapt to each other’s changing behaviors. The Self Critic helps the agent optimize its individual performance, while the Other Critic accounts for the impact of other agents’ actions. This dual-critic approach enables more effective learning in complex multi-agent environments where both individual and collective behaviors are crucial.

## 5 Experiments

### Iterated Games

We first review the two iterated games, specifically the the Iterated Prisoner’s Dilemma (IPD) and explain how these games can be modeled as a two-agent Markov Decision Process (MDP).

Table 1 shows the per-step payoff matrix of the prisoners’ dilemma. In a single-shot prisoners’ dilemma, there is only one Nash equilibrium, where both agents defect. In the infinitely iterated prisoners’ dilemma, the folk theorem shows that there are infinitely many Nash equilibria. Two notable ones are the always defect strategy (DD) and tit-for-tat (TFT). In TFT, each agent starts with cooperation and then repeats the previous action of the opponent. The average returns per step in self-play are -1 and -2 for TFT and DD, respectively.

**Table 1:**
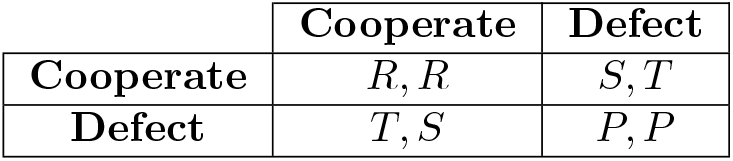
Prisoner’s Dilemma Payoff Matrix

|  | **Cooperate** | **Defect** |
| --- | --- | --- |
| **Cooperate** | $R, R$ | $S, T$ |
| **Defect** | $T, S$ | $P, P$ |

### Environment Description

The Iterated Prisoner’s Dilemma can be modeled as a Markov Decision Process (MDP), defined by the tuple (*S, A*, *T*, ℛ, *γ*). The state space *S* = {(*a*_1_, *b*_1_, …, *a*_*k*_, *b*_*k*_) | *a*_*i*_, *b*_*i*_ ∈ {0, 1}, *i* = 1, …, *k* represents the actions of both players over the past *k* rounds, where 0 is Cooperate and 1 is Defect, and *k* is the memory length. The action space *A* = {0, 1} corresponds to the same choices.

The transition function *T*: *S*× *A*× *S* → [0, 1] gives the probability of transitioning between states given actions, which is deterministic in this fully observable setting. The reward function ℛ: *S* ×*A* →ℝ is based on the standard Prisoner’s Dilemma payoff matrix, with *T > R > P > S*, where *T* is the temptation to defect, *R* is the reward for mutual cooperation, *P* is the punishment for mutual defection, and *S* is the sucker’s payoff.

The discount factor *γ* ∈ [0, 1] represents the importance of future rewards. In this framework, an agent’s strategy *π* : *S* →*A* maps the current state (fully observed history) to actions. We tested this model for memory lengths of 1, 2, and 3, resulting in |*S*| = 4^*k*^ states for memory length *k*. This MDP formulation allows for the exploration of dynamic strategies that can adapt based on the fully observable history of both players’ actions.

### Dataset

We utilized datasets derived from both human experiments and animal models to provide a comprehensive view of strategic decision-making in social dilemmas.

### Human Iterated Prisoner’s Dilemma Dataset

^1^The primary dataset originates from the work of Montero-Porras et al., encompassing experimental data from long Iterated Prisoner’s Dilemma (IPD) experiments. The data were collected at the Brussels Experimental Economics Laboratory (BEEL) of the Vrije Universiteit Brussel (VUB) and are available on Zenodo. The study includes two treatments with fixed and shuffled partners, involving a total of 188 participants across 12 sessions. Participants played the IPD on isolated laptops to prevent communication, and the game parameters were set as *R* = 3, *T* = 4, *S* = 0, and *P* = 1. Table 2 presents the payoff matrix containing the per-round rewards used in the two treatments. In both treatments, participants could observe their partner’s previous action, even when the partner changed from the previous to the current round.

**Table 2:** Payoff matrix of prisoners’ dilemma.

|  | C | D |
| --- | --- | --- |
| C | (3, 3) | (0, 4) |
| D | (4, 0) | (1, 1) |

### Training Details

We trained reinforcement learning (RL) agents to investigate the strategies that emerge in the Iterated Prisoner’s Dilemma (IPD). Each round of the IPD consists of the same type of agents playing against each other simultaneously. In each iteration, agents must choose to either cooperate or defect. Importantly, each agent has full access to the reward and state information of the other player, creating a fully observable environment. We tested a spectrum of RL approaches, including Deep Q-Network (DQN; Mnih et al., 2015) as our value-based method, Proximal Policy Optimization (PPO; Schulman et al., 2017) as our policy-based method, and Advantage Actor-Critic (A2C; Mnih et al., 2016) as our actor-critic method. Additionally, we introduced a new variant, the Theory of Mind Actor-Critic (TOMAC), which was described in the Methods section.

For all algorithms, we used fully connected neural networks with one-hot encodings of state representations as input. This approach allows for efficient representation of discrete states and facilitates learning in the IPD environment. Hyperparameter sweeps were performed to optimize the learning process. We tested network architectures with 256, 64, and 32 hidden units. Learning rates were set at 1e-07, 0.001, and 0.3, while discount factors (*γ*) were 0.5, 0.75, and 0.9999. Exploration was implemented using an *ϵ*-greedy algorithm with *ϵ* set to 0.05. We also included a learning rate for the temperature parameter, *lr*_*T*_. For each hyperparameter configuration, the agents were trained for 20 iterations. Each iteration consisted of 100 games (rounds) of the IPD. Agents received observations based on the context length, which was set to 1, 2, or 3 (i.e., previous step, previous two steps, or previous 3 steps) to align with biological plausibility. This setup allowed us to explore a range of learning parameters and network architectures while maintaining a biologically plausible observation space.

### Evaluation Metrics

To assess the performance of the algorithms and compare them with human behavior, we employed several key metrics:

#### Cooperation Rate

The cooperation rate is a crucial measure of mutual cooperation, reflecting the emergence of successful strategies in the Iterated Prisoner’s Dilemma (IPD). This metric is particularly significant as the winning strategy for long IPD games is typically Tit-for-Tat, which often results in both players cooperating. The cooperation rate is calculated as a running average: at each step, if both agents cooperate, it scores 1; otherwise, it scores 0. This running average provides a dynamic measure of how often agents achieve mutual cooperation over time, serving as both an indicator of optimal strategy emergence and a standard measure in behavioral modeling for comparing human data and models (Nay and Vorobeychik 2016).

#### Cumulative Reward

We track the cumulative reward, which is a running average of the collective reward achieved by both agents at each time step, divided by 2 to represent the average reward per agent. Given our IPD reward structure (T=4, S=0, P=1, R=3), the maximal achievable cumulative reward over 100 rounds is 300, obtained through consistent mutual cooperation. This cumulative reward metric offers insights into the average performance of the agents in both cooperative and competitive settings, allowing us to assess how well the algorithms maximize long-term rewards.

#### Similarity Score

We utilized a similarity score to quantify how closely the algorithms’ decisions matched human decisions despite facing different opponent moves. This similarity score is computed as a running average of stepwise comparisons of actions: at each step, if the algorithm’s action matches the human action, it scores 1; otherwise, it scores 0. The net similarity score achieved at the 100th step is considered the final similarity score measure, providing a comprehensive evaluation of the algorithm’s alignment with human behavior over the entire interaction sequence. This score is crucial for assessing the ecological validity of our models and identifying potential discrepancies between AI strategies and human decision-making processes.

### Experimental Design

We conducted two types of experiments: online learning and constrained offline imitation learning experiments. In the online learning experiments, we implemented the RL algorithms in the IPD environment in a standard online fashion, allowing the agents to learn from their interactions without restrictions. This approach aligns with traditional reinforcement learning methods, where agents update their policies based on continuous feedback from the environment [22].

In the constrained offline imitation learning experiments, we employed a methodology similar to offline imitation learning or behavioral cloning [8, 16]. We used the datasets mentioned above to control the observations and rewards that the algorithm receives. Specifically, we constrained the agent’s experience by feeding it the same observations and rewards that the humans experienced, but we also recorded the agent’s preferred actions under these constraints. This method allows the agent to learn from a fixed dataset without additional interactions with the environment, mirroring the offline imitation learning paradigm.

As illustrated in Figure 2, for the human data comparison, we tested pairs of the same type of agents playing against each other, which will be referred to as “Agent vs. Agent” in the results section. The figure depicts human participants playing the IPD, highlighting context lengths of 1, 2, and 3 in the game. This setup allows us to assess how well the RL agents replicate human decision-making patterns in the IPD when interacting with similar counterparts.

**Figure 1.**
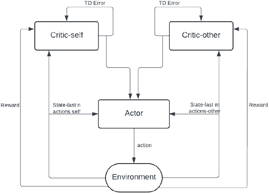
Figurative description of the TOMAC.

**Figure 2.**
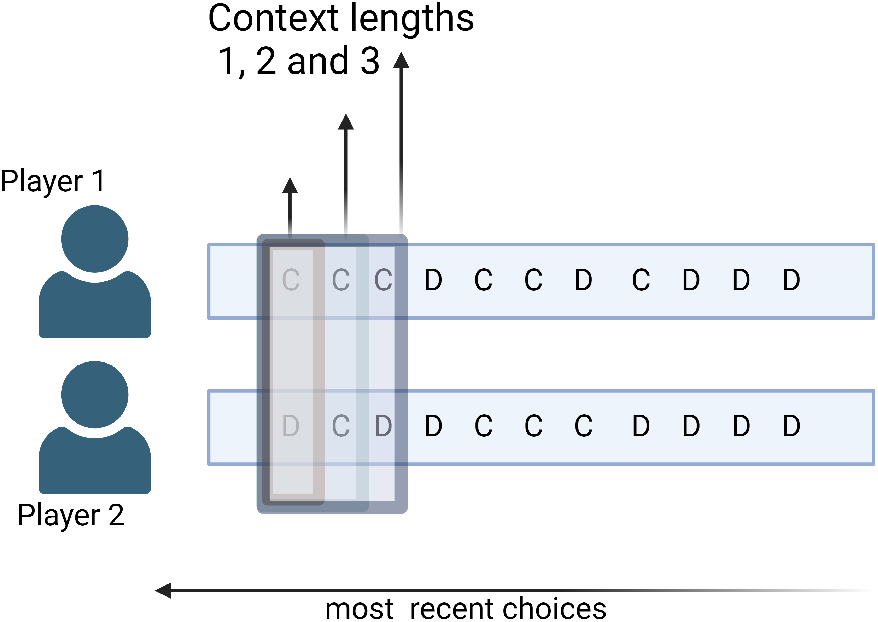
Illustration of human participants playing the IPD .Context lengths 1,2,3 in the contet of the IPD are depicted in the figure

**Figure 3.**
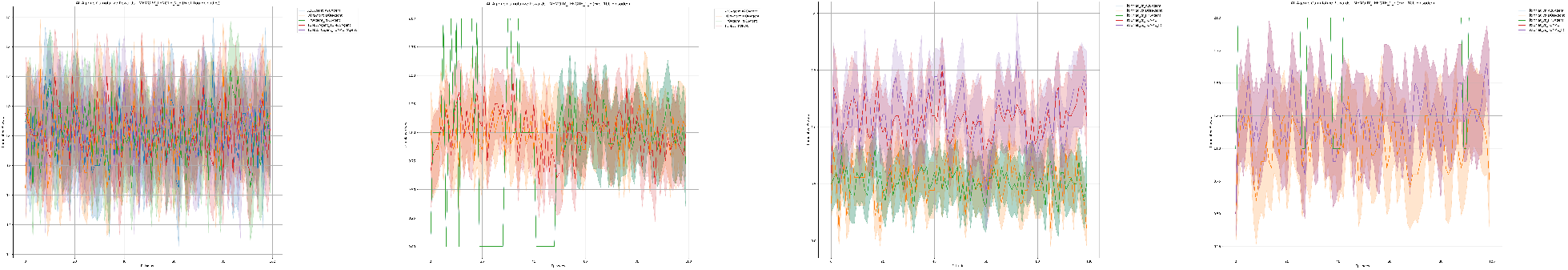
Average rewards: (a) Agents vs Agents (Unconstrained), (b) Agents vs Agents (Constrained), (c) TFT vs Agents (Unconstrained), (d) TFT vs Agents (Constrained). TOMAC and TOMAC with internal reward achieved higher average reward for TFT in the unconstrained scenario (c), while PPO achieved higher rewards in the constrained scenario (d), similar to its performance in the constrained agent vs agent scenario (b). There is no clear distinction in the average rewards for the unconstrained agent vs agent scenario (a).

For the human data, we further divided the participants into clusters based on their behaviors. Participants were clustered based on whether they converged to cooperation or defection in the last 15 trials out of 100, and whether they exhibited high or low variability in their actions. The clusters are: converged to cooperation (high initial variability), converged to defection (high initial variability), converged to cooperation (low variability), converged to defection (low variability), and unconstrained learning. By dividing the data in this manner, we aimed to capture different strategic behaviors exhibited by the participants. We then compared the algorithms’ performance to the human data within each cluster to assess how well the RL agents could replicate the diverse human strategies.

The objective was to observe the divergence in policy when the input to the artificial agents is the same as that of the humans. By comparing the agents’ preferred policies to the actual ones, we aimed to identify differences in strategic decision-making processes. All evaluation metrics were calculated using the preferred policies and compared to the actual ones.

## 6 Results

### 6.1 Cumulative Reward Analysis

#### 6.1.1 Agent vs. Agent

We analyzed agent performance in three scenarios: online learning, constrained online learning with cooperative strategies, and constrained online learning with defective strategies. In the online learning scenario, we observe significant differentiation between algorithms. A2C demonstrates superior performance, achieving a cumulative reward of 180, followed by TOMAC at 150. DQN and PPO show similar, but lower, performance at 110 and 104 respectively. In the constrained online learning scenario with cooperative strategies, all AI agents show similar performance, with cumulative rewards ranging from 95 to 103. TOMAC slightly leads at 103, followed closely by A2C at 101. However, human performance significantly outperforms all AI agents in this scenario, reaching approximately 155 before the final drop. This substantial performance gap suggests that AI algorithms struggle to capture or imitate cooperative strategies compared to humans even when constrained to them. In the constrained online learning scenario where we use defective human strategies, we observe an interesting reversal. AI agents perform similarly to each other and to the human defectors, with cumulative rewards ranging from 110 to 116, led by A2C at 116. This suggests that AI agents might be better at adapting to or exploiting defective behavior than humans, when constrained by human-like strategies.

A key observation across scenarios is the reduction in performance variance in constrained online learning compared to the online learning scenario. This variance reduction suggests that the agents effectively learn from and replicate human behavior patterns in constrained online learning, though it also implies a potential limitation in discovering novel or superior strategies, which might explain the lower cumulative rewards in the constrained cooperative scenario. Detailed tables of cumulative rewards, including breakdowns by memory length, are provided in the supplementary material.

#### 6.1.2 Agent vs. Tit-for-Tat (TFT)

Our comparative analysis of agent performance in the Iterated Prisoner’s Dilemma (IPD) revealed significant variations across agent types and memory lengths. AI agents’ performance varied markedly based on memory length. For memory length 1, AI agents showed consistent performance, scoring between 105–120, with A2C slightly outperforming others at approximately 120. However, memory length 2 introduced significant performance disparities: TOMAC and TOMAC with internal reward excelled (110–115), while A2C, DQN, and PPO underperformed (50–55). This divergence suggests that increased memory length enables more complex strategies, benefiting some algorithms while challenging others. These findings underscore the potential for cross-disciplinary research between neuroscience and artificial intelligence to enhance the adaptability and performance of reinforcement learning algorithms in complex, strategic environments.

**Note:** Detailed figures illustrating these performance comparisons can be found in the supplementary material.

### 6.2 Evaluation of Similarity Scores in Constrained Learning

#### 6.2.1 Agent vs. Agent

Our findings indicate that **TOMAC** is the algorithm that most closely replicates human behavior, particularly under high variability conditions:

- In the *Mutual Cooperation (High Variability)* condition, TOMAC achieved similarity scores around 63%, outperforming other algorithms under the same conditions. This reflects the complex and adaptive nature of human cooperative behavior in unpredictable environments.
- In the *Mutual Defection (High Variability)* scenario, TOMAC maintained moderate similarity scores between 62% and 64%. These results suggest that TOMAC captures the fluctuating patterns characteristic of human defection strategies in uncertain settings.

#### 6.2.2 Agent vs. TFT

Conversely, **A2C** and **PPO** appear to better replicate animal behavior, especially under low variability conditions:

- In the *Mutual Cooperation (Low Variability)* scenario, both A2C and PPO achieved high similarity scores of approximately 88% across all evaluation metrics. This suggests a strong alignment with animal cooperative strategies in stable environments, where behavior tends to be more predictable.
- In the *Mutual Defection (Low Variability)* setting, A2C and PPO maintained high similarity scores around 77%, indicating effective replication of animal defection strategies when environmental factors are consistent.

### 6.3 Cooperation Rate Analysis

#### 6.3.1 Agent vs. Agent

We observe distinct cooperation patterns among AI agents across constrained and unconstrained experiments. In constrained scenarios, where agents’ actions are influenced by human data, PPO and A2C exhibit similar behaviors, showing high sensitivity to initial conditions and variance levels. They maintain high initial cooperation rates across all scenarios but display more pronounced fluctuations in defector scenarios (Figures 4a–d). This suggests these algorithms adapt dynamically to environmental changes, potentially at the cost of stability.

**Figure 4.**
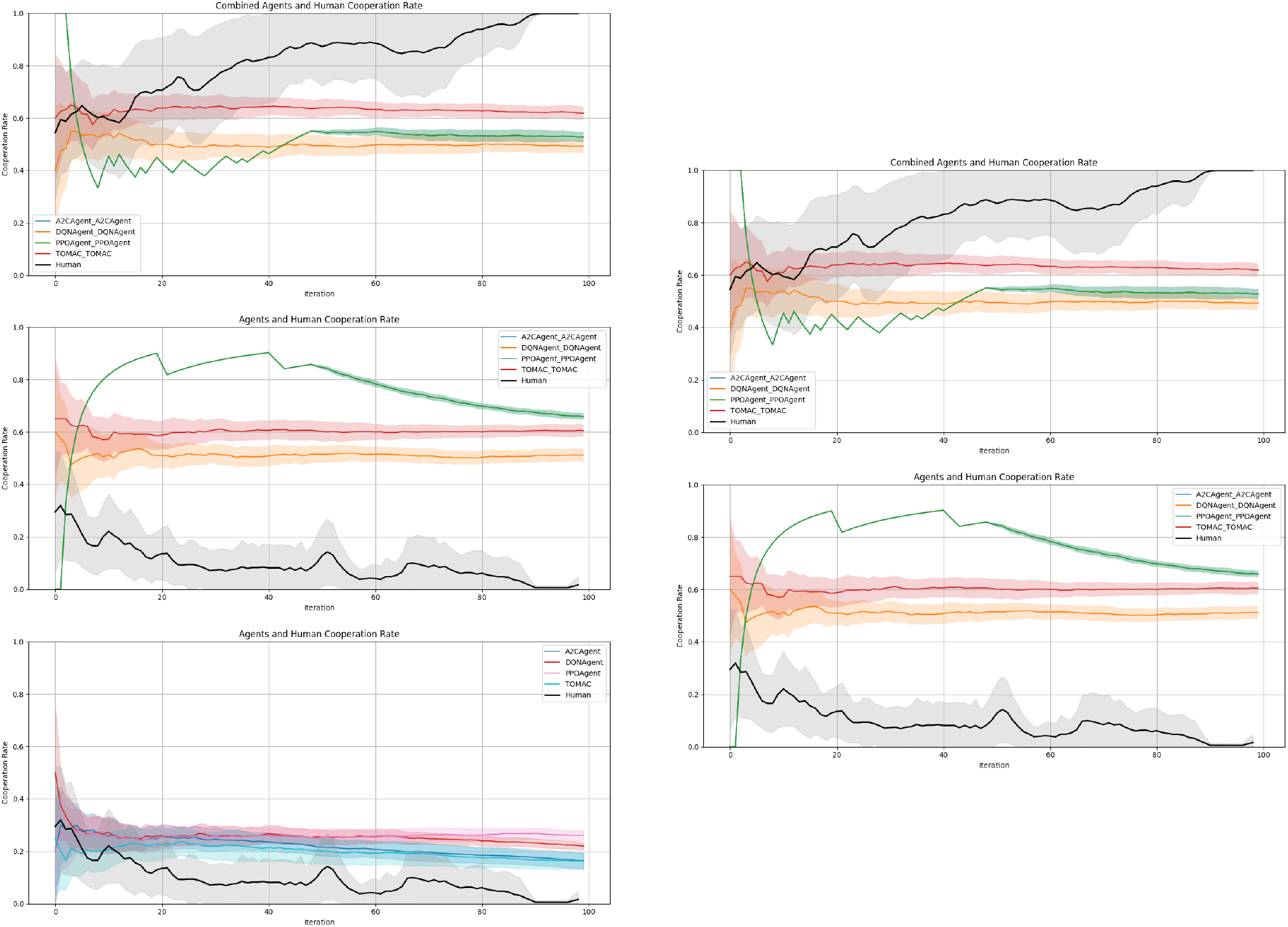
Cooperation rates across different learning scenarios in Agent vs. Agent experiments. (a) Constrained learning using cooperation (high initial variability) (b) Constrained learning using defection (high initial variability) (c) Constrained learning using cooperation (low variability) (e) Constrained learning using defection (low variability) (d) Unconstrained learning.

In constrained scenarios (a–d), PPO and A2C show high sensitivity to initial conditions and variance levels, while TOMAC maintains consistent behavior around 60% cooperation. DQN displays stable performance at 50% cooperation. All AI agents show higher cooperation than humans in defector scenarios. In the unconstrained scenario (e), AI agents maintain higher cooperation rates (25–30%) compared to human defectors, whose rates decrease to nearly 0%.

#### Constrained Experiments

In constrained scenarios, we observe distinct patterns among AI agents. PPO and A2C exhibit high sensitivity to initial conditions and variance levels, maintaining high initial cooperation rates but showing pronounced fluctuations, especially in defector scenarios. In contrast, TOMAC demonstrates the most consistent behavior, maintaining a steady cooperation rate slightly above 60% across all scenarios, indicating a robust, balanced strategy. DQN displays stable performance, consistently maintaining around 50% cooperation with low sensitivity to changing conditions.

Notably, all AI agents show higher cooperation rates than humans in defector scenarios, suggesting a robust bias towards cooperation. Agent behaviors vary based on the variability of human strategies. High-variance scenarios lead to more fluctuations in agent behavior, particularly for PPO and A2C, while low-variance scenarios result in more stable behaviors across all agent types.

#### Unconstrained Experiments

In unconstrained scenarios, where agents are not influenced by human data, we observe a distinct pattern. All AI agents maintain significantly higher cooperation rates (25–30%) compared to human defectors throughout the iterations. In contrast, human defector cooperation rates start low (around 30%) and decrease to nearly 0% by the end. This suggests that when unconstrained by human-derived strategies, AI agents develop more persistently cooperative approaches.

The consistency in higher cooperation rates among AI agents, especially in defector scenarios, may indicate either an ability to maintain cooperation or a potential limitation in adapting to highly competitive environments. These findings highlight the significant impact of learning conditions (constrained vs. unconstrained) and human strategy variability on AI agent behavior and cooperation rates in the Iterated Prisoner’s Dilemma.

### 6.4 Agent vs. Tit-for-Tat (TFT)

Figure 7. Cooperation rates of agents against Tit-for-Tat strategy in constrained and unconstrained experiments. (a) Constrained scenario: Most agents show stable cooperation rates with narrow confidence intervals. A2C and DQN trend towards stable cooperation after initial fluctuations. PPO maintains high cooperation throughout. TOMAC and TOMAC with Internal Reward show lower cooperation rates. (b) Unconstrained scenario: Higher variability in cooperation rates with wider confidence intervals. A2C has significant initial fluctuations but stabilizes over time. PPO and DQN start with higher cooperation but decrease, suggesting adaptive strategies shifting towards defection. TOMAC and TOMAC with Internal Reward consistently show lower cooperation rates in both scenarios. The consistent lower cooperation rates of TOMAC in both scenarios suggest a preference for defection regardless of constraints.

#### 6.4.1 Constrained Experiments

The constrained experiments (Figure 5) show stabilization in cooperation rates for most agents, with narrow confidence intervals indicating consistent behavior. Tit-for-Tat vs. A2C and Tit-for-Tat vs. DQN display a trend towards stable cooperation after initial fluctuations, whereas Tit-for-Tat vs. PPO maintains high cooperation with minimal variance throughout. Lower rates observed in Tit-for-Tat vs. TOMAC and Tit-for-Tat vs. TOMAC with Internal Reward suggest less cooperative strategies within clamped settings.

**Figure 5.**
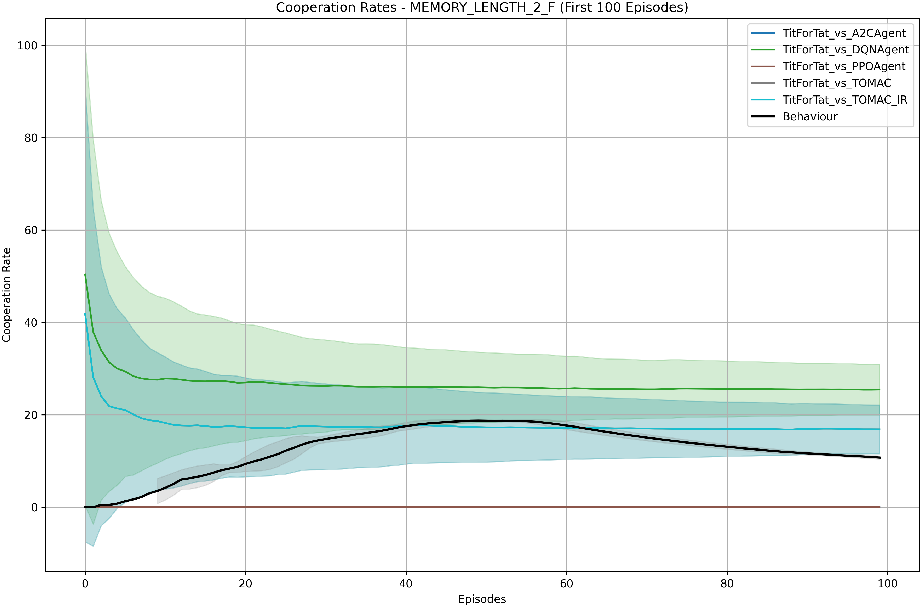
Constrained Cooperation Rates

#### 6.4.2 Unconstrained Experiments

The unconstrained experiments (Figure 6) reveal higher variability in cooperation rates across agents, with wider confidence intervals signifying less consistent behaviors. Significant initial fluctuations in Tit-for-Tat vs. A2C stabilize over time. Tit-for-Tat vs. PPO and Tit-for-Tat vs. TOMAC with Internal Reward start with higher cooperation but tend to decrease, indicating adaptive strategies that may shift towards defection. Similar to clamped experiments, Tit-for-Tat vs. TOMAC and Tit-for-Tat vs. TOMAC with Internal Reward consistently show lower cooperation rates, reinforcing their preference for defection in less restricted settings.

**Figure 6.**
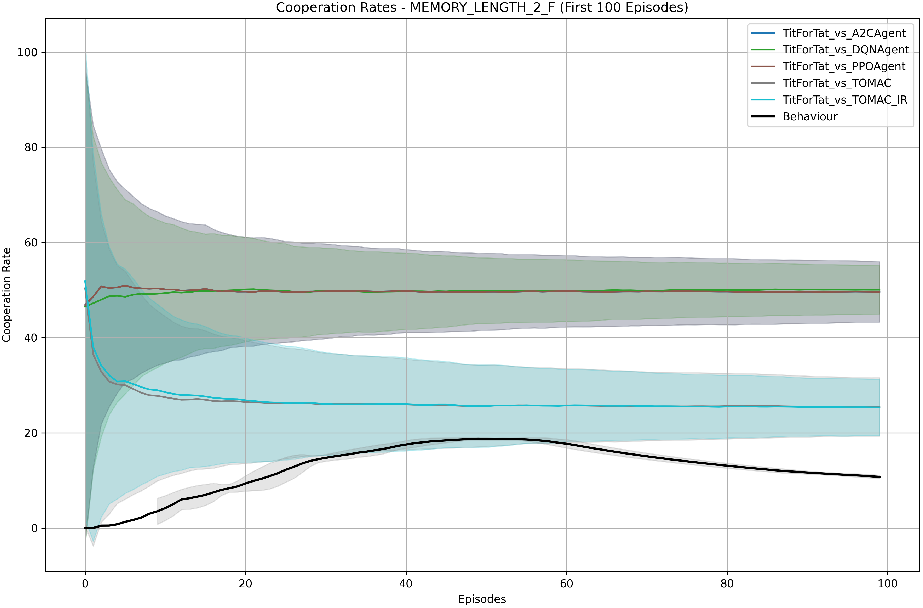
Unconstrained Cooperation Rates

## 7 Conclusion

Our analysis of cooperation rates and similarity scores in the Iterated Prisoner’s Dilemma reveals crucial insights into the decision-making processes of humans and AI agents. The data suggests a clear distinction in the learning capabilities and strategies employed by humans compared to those of current AI algorithms.

Humans demonstrated an ability to learn the rules of the game and adopt increasingly effective strategies as the memory length increased. This is evidenced by the increasing similarity scores for human behavior as the context window expands, particularly aligning with TOMAC’s performance. This suggests that humans are likely using a more complex algorithm, similar to TOMAC, which benefits from a larger context window, allowing for more sophisticated strategy formulation and adaptation.

AI agents, particularly TOMAC, showed consistent performance across different memory lengths, with slight improvements as the context window increased. Although other AI methods such as A2C and PPO replicate certain aspects of human decision-making in specific scenarios, they do not fully capture the adaptive and flexible strategies exhibited by humans in extended gameplay.

These findings highlight the significant differences in strategic decision-making capabilities between humans and AI. Humans appear to possess the most flexible and adaptive approach, capable of leveraging increased historical information to improve their strategies, while AI agents still lag behind in long-term strategic adaptation.

## 8 Limitations and Future Work

While our study provides valuable insights into strategic decision-making in biological and artificial agents, several limitations warrant consideration and point to directions for future research.

A primary limitation of our current TOMAC model is its assumption of a pre-decomposed value function in fully observable environments. This assumption may not hold in more complex, partially observable scenarios. Future work will need to address this limitation by incorporating a Value Decomposition Network (VDN) [18] or similar approaches at the head of the network. This addition will allow TOMAC to automatically decompose the value function into self and other components, making it more adaptable to a wider range of environmental complexities and observability conditions.

Another significant limitation is TOMAC’s current focus on two-player scenarios. Future work will aim to scale this model to multi-agent systems, incorporating multiple critics to reflect varying group sizes. This expansion presents significant computational challenges and will require exploring hierarchical modeling approaches [14]. To validate these developments, we plan to conduct human experiments with increasing numbers of players, which will help reveal the limitations of human strategic decision-making as group size grows.

Although we have benchmarked TOMAC against baseline algorithms, further comparisons with a broader range of opponent modeling techniques are necessary. Future studies will examine TOMAC alongside Fictitious Play [2], Bayesian Strategy Inference, and Recursive Modeling [6]. We also plan to compare TOMAC with gradient-based multi-agent learning approaches like LOLA [5] and POLA [9] in more complex environments beyond two-player games. These comparisons will provide a more comprehensive assessment of TOMAC’s robustness and capabilities.

The current implementation of TOMAC, while inspired by neuroscientific findings, still simplifies many aspects of biological decision-making processes. To enhance the model’s biological plausibility and performance in longer games, we aim to integrate more sophisticated memory mechanisms. These include adaptive memory decay, episodic memory, and working memory, which could significantly improve the model’s ability to handle extended gameplay scenarios [3].

A significant avenue for future work lies in strengthening the neurobiological grounding of our model. We plan to integrate neuroimaging and neural recording data to refine TOMAC’s biological plausibility and investigate how different brain regions contribute to strategic decision-making, particularly in larger group settings [10].

By addressing these limitations, we aim to create a more comprehensive, biologically grounded model of strategic social decision-making. Such a model has the potential to offer deeper insights into human behavior, contribute to evolutionary game theory, and inform the design of sophisticated artificial agents capable of navigating complex social environments.

## Footnotes

1 This dataset is from [13] and is available at [12].

